# Production of human translation-competent lysates using dual centrifugation

**DOI:** 10.1101/2021.05.27.445927

**Authors:** Lukas-Adrian Gurzeler, Jana Ziegelmüller, Oliver Mühlemann, Evangelos D. Karousis

## Abstract

Protein synthesis is a central process in gene expression and the development of efficient *in vitro* translation systems has been the focus of scientific efforts for decades. The production of translation-competent lysates originating from human cells or tissues remains challenging, mainly due to the variability of cell lysis conditions. Here we present a robust and fast method based on dual centrifugation that allows for detergent-free cell lysis under controlled mechanical forces. We optimized the lysate preparation to yield cytoplasm-enriched extracts from human cells that efficiently translate mRNAs in a cap-dependent as well as in an IRES-mediated way. Reduction of the phosphorylation state of eIF2α using recombinant GADD34 and 2-aminopurine considerably boosts the protein output, reinforcing the potential of this method to produce recombinant proteins from human lysates.

## INTRODUCTION

Cell-free translation is the process of synthesizing proteins in extracts without the use of intact cells. During the last five decades, cell-free extracts have been used to recapitulate key steps of translation and have provided valuable insights into this fundamental process (1). Bacterial-based lysates have been extensively used to develop, understand, and expand the applications of *in vitro* translation (2). Ranging from the historical experiments that led to the determination of the genetic code (3,4) to recent advances that allow the design of orthogonal *in vitro* translation systems (5,6), cell-free protein synthesis has revolutionized the possibilities of synthetic biology by employing and expanding the properties of *in vitro* translation. Moreover, cell-free systems have allowed the production of numerous recombinant proteins with clear advantages compared to production in living cells. The possibility of genetic manipulation, the incorporation of unnatural amino acids and the tolerance to toxic proteins in cell-free extracts have extended the limits of synthetic biology significantly (2).

Even though the production and use of human lysates is of particular interest, it has not been equally developed. Human cell-derived lysates provide a better environment for the proper folding of human proteins as they facilitate the incorporation of human-specific post-translational modifications. Moreover, they are an appealing system to study translation-related processes in small scale or high throughput settings. However, several key constraints like the cost and technical difficulties in the reproducible production of eukaryotic translation systems compared to prokaryotic have limited their industry and research usage (7).

The relatively low and variable protein synthesis output are the most important reasons that impede the routine application of human *in vitro* translation systems (1). To date, the only commercially available mammalian *in vitro* translation system that allows for cap-dependent translation is the rabbit reticulocyte lysate (RRL) (8). In contrast, the currently commercially available human *in vitro* translation systems depend on Internal Ribosome Entry Site (IRES)-mediated initiation and perform poorly on normal ^7^mG-capped mRNAs (Thermo Fisher Scientific, Cat No. 88890). The RRL system has proven to be a valuable source to study several aspects of mammalian translation. However, it is accompanied by unsurpassed caveats due to reticulocytes being special cells regarding their translation activity and regulation. The requirement of complex manipulation of animal tissue, the increased concentration of endogenous globin mRNAs, and the abundance of RNase M and other tissue-specific nucleases (9) are substantial limitations of this system. Moreover, RRL cannot be genetically modified to deplete or enrich certain protein factors (10). Previous efforts have advanced the use of mammalian cells to produce translation-competent lysates. Chinese hamster ovary cells (CHO) have been used to develop an *in vitro* translation system that allows for IRES-dependent translation of reporter constructs and has been shown to produce functional glycosylated proteins of interest (11). Lysates of human origin that support cap-dependent and independent *in vitro* translation of reporter mRNAs originate from HeLa cells (12–14) or from HEK-293 cells for the study of native mRNP complexes *in vitro* (15). However, issues concerning the variability of lysis parameters, such as the applied mechanical force, have not been satisfactorily solved.

In our previous efforts to address mechanistic questions about translation termination, we developed an *in vitro* translation system based on HeLa cells that was used to monitor ribosomal density at the termination codon of reporter mRNAs (16). This protocol was optimized to prepare lysates using the shearing forces of syringe treatment under hypotonic conditions to break up the cells and release functionally active molecules that can reconstitute the complex translation process *in vitro* (16), based on a previously reported protocol (14). During this development, we observed that cell lysis is the most crucial step for the success of our *in vitro* system. To further improve the efficiency and the reproducibility of this step, we applied dual centrifugation (DC) as a novel approach to lyse cells. Dual centrifugation differs from regular centrifugation by combining the simultaneous rotation around two axes. As a consequence, the centrifugal acceleration constantly changes, resulting in powerful agitation of the sample material inside the vials (Fig. 1A). Thus, in contrast to conventional centrifugation that separates components, DC leads to efficient sample homogenization (17,18). Adapting this technique as a cell lysis method, we streamlined the production of translation-competent lysates in various batch sizes in a reproducible manner. The resulting lysates allowed us to dissect the role of Nsp1, a potent virulent factor produced during the early steps of infection by SARS-CoV-2 in human cells (19).

**Figure 1:**
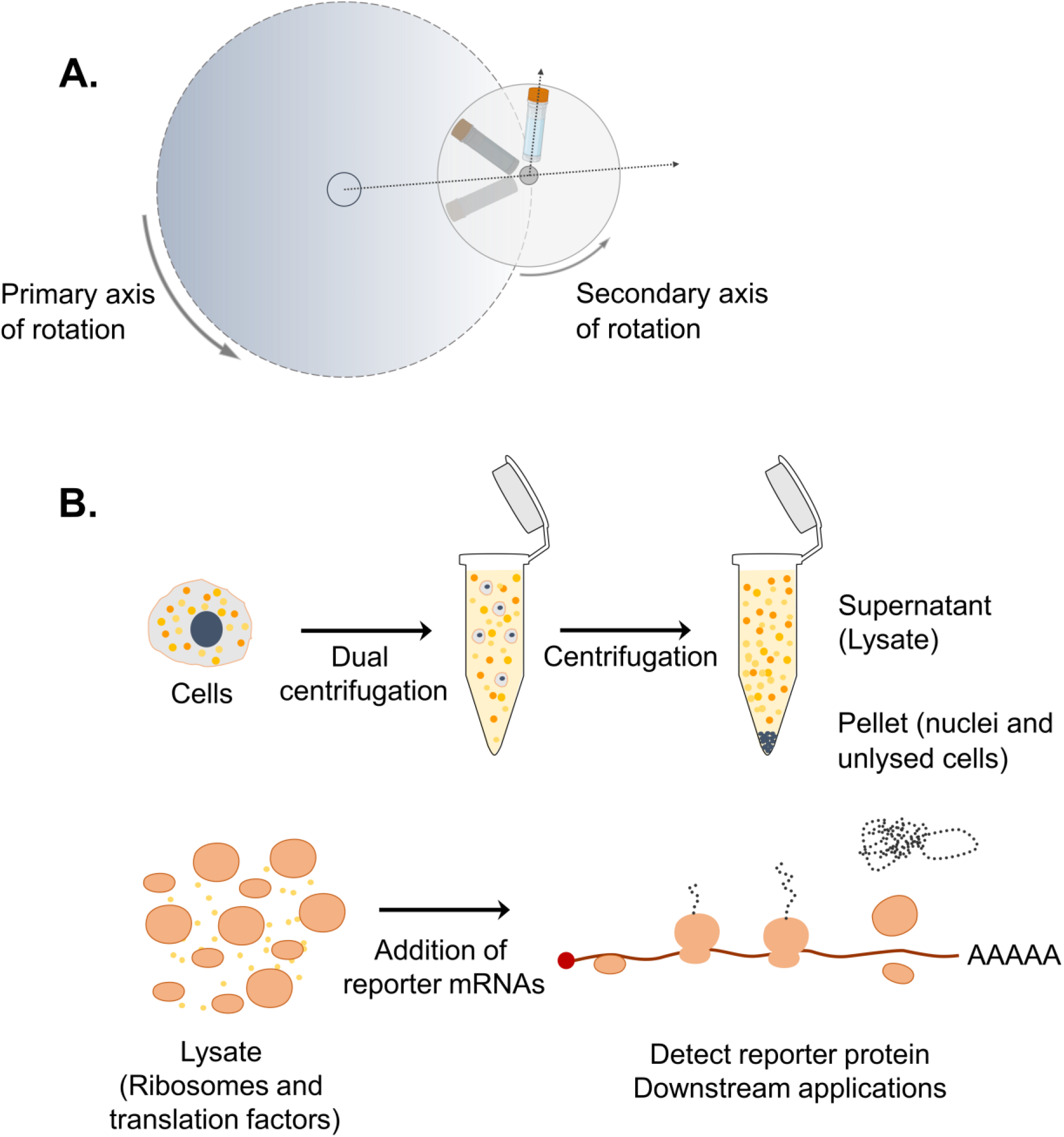
Dual centrifugation as a cell lysis method to produce translation-competent lysates. **A**. Schematic representation of DC. During DC, the samples rotate simultaneously around two axes of symmetry. Dashed arrows indicate the direction of centrifugal acceleration. **B**. Graphical overview of the steps for the production of translation-competent lysates using DC.

Here we document that this new, improved protocol using dual centrifugation enables the efficient and reproducible production of translation-competent cytoplasmic cell extracts from HeLa S3 cells in large amounts. We show that dual centrifugation can be used for routine cell lysis under different buffer compositions, and we investigate the conditions that give rise to lysates that translate reporter mRNAs in a cap- and IRES-dependent manner. We perform biochemical characterization of the lysates, provide technical details for their production, and show that by modifying the phosphorylation state of eIF2α we can increase protein synthesis considerably.

## MATERIAL AND METHODS

### Cell lines and growth conditions

HeLa S3 cells (Sigma-Aldrich, ECACC, Cat. No. 87110901) were cultured in DMEM/F-12 (Gibco, Cat. No. 32500-043) containing 10% FCS, 100 U/ml penicillin and 100 μg/ml streptomycin (termed DMEM+/+) at 37 °C and 5% CO_2_ in humid atmosphere. To start a new culture from a cryopreserved stock (stored in 50% FCS, 40% DMEM +/+ and 10% DMSO), 1-2×10^6^ cells were seeded into a 75 cm^2^ flask (TPP, Cat. No. 90026). After reaching a confluency of approximately 80% as adherent cells, they were trypsinized and transferred into a suspension culture. The suspension cultures were then grown in volumes between 20 ml and 500 ml under constant agitation (Corning flasks, Cat. No. 431405, 431147 and 431255) within a range of 0.2×10^6^ -1×10^6^ cells/ml.

### Preparation of translation-competent lysates

Lysates were prepared from non-synchronized HeLa S3 cell cultures grown to a cell density ranging from 0.8×10^6^ to 1×10^6^ cells/ml. Cells were pelleted (200 *g*, 4 °C for 5 min), washed twice with cold 1x PBS pH 7.4 and resuspended in cold hypotonic lysis buffer (10 mM HEPES pH 7.3, 10 mM K-acetate, 500 μM MgCl_2_, and 1x protease inhibitor cocktail (Bimake, Cat. No. B14002)) or translation-competent (lysis) buffer (33.78 mM HEPES pH 7.3, 63.06 mM K-acetate, 0.68 mM MgCl_2_, 54.05 mM KCl, 0.05 mM amino acid mixture, unless otherwise stated (Promega, Cat. No. L4461), 13.51 mM Creatine Phosphate, 229.73 ng/ml Creatine kinase and 1x protease inhibitor cocktail (Bimake, Cat. No. B14002)) at a final concentration of 2×10^8^ cells/ml in 2 ml screw cap microtubes (Sarstedt, Cat. No. 72.693). To assess the translation activity of lysates from frozen cells, cell pellets were stored at -80 °C overnight before resuspension in lysis buffer. Cells were lysed by dual centrifugation at 500 RPM (unless otherwise stated) at −5 °C for 4 min using ZentriMix 380 R system (Hettich AG) with a 3206 rotor with 3209 adapters. Cell lysis was monitored by staining a 1:10 (f.c.) diluted sample before and after dual centrifugation with 0.2% (f.c.) Trypan Blue (Thermo Fisher Scientific, Cat. No. T10282) using a light microscope with EVE Cell counting slides (NanoEnTek, Cat. No. EVS-050). The lysate was centrifuged at 13,000 *g*, 4 °C for 10 min and the supernatant (the lysate) was aliquoted, snap-frozen and stored at -80 °C. The lysate could be thawed and refrozen multiple times for subsequent applications with only a minor loss of translation efficiency.

### Plasmids and cloning

The DNA template sequences of the different reporters used in this study are depicted in sup. Fig. 3.The p200-6xMS2 plasmid encoding the RLuc reporter used for *in vitro* transcription is described in (19). We created the dual-luciferase reporter pCRIIRLuc-UAA-FLucA80 by ligating *Not*I-*Kpn*I digested pCRII-hRLuc-200bp-A80 (16) and the PCR-amplified Luc ORFs from the plasmid p2luc-UAA (20) (digested with the same restriction enzymes). The ECMV IRES sequence was subcloned into pCRII-RLuc-UAA-FLuc-A80 from the pT7CFE1-CHis plasmid (Thermo Fisher Scientific, Cat. No. 88860) by In-Fusion cloning (Takara, Cat. No. 639650) to create pCRII-RLuc-IRES-FLuc-A80. A second *Hind*III restriction site was deleted by site-directed mutagenesis because this enzyme is used for linearization before run-off *in vitro* transcription. To clone pCRII_SC-LD_SMG6-3xFLAG_A80, the SMG6 coding sequence (CDS) along with a C-terminal 3xFLAG was cloned into pCRII-FL-hRLuc described in (19) by removing the hRLuc sequence and only leaving the SARS-CoV-2 Leader sequence (SC-LD) of the SARS-CoV-2 5’UTR using InFusion cloning. To obtain pCRII_SC-LD_3xFLAG-HBB-CTE_A80, the beta globin CDS (HBB) with a 70-nucleotide long C-terminal extension (CTE) was cloned into the pCRII_SC-LD_SMG6-3xFLAG_A80, by replacing the SMG6 coding sequence and moving the 3xFLAG tag to the N-terminus using InFusion cloning.

### *In vitro* transcription of reporter mRNAs

Plasmids used for *in vitro* transcription were linearized using a restriction site located downstream of the poly(A) sequence in a reaction containing 40 ng/μl DNA (4 μg total) and 0.5 U/μl restriction enzyme (*Hind*III-HF; NEB, Cat. No. R3104L for pCRII-hRLuc-200bp-A80, pCRII-p2luc-UAA-A80 and pCRII_SC-LD_3xFLAG-HBB, and *Cla*I; NEB, Cat. No. R0197S for pCRII_SC-LD_SMG6-3xFLAG_A80) in 1x CutSmart buffer (NEB, Cat. No. B7204S). The digest was monitored by running 200 ng of the DNA on a 1% agarose gel in 1x TAE. The rest of the reaction was purified using the MACHEREY-NAGEL NucleoSpin® Gel and PCR Clean-up kit (Cat. No. 740609.50). The isolated DNA was then quantified by measuring OD_260_.

Using the linearized plasmid as a template (up to 4 μg in total and 40 ng/μl f.c.), the reporter mRNA was *in vitro* transcribed in a reaction containing 1x OPTIZYME transcription buffer (Thermo Fisher Scientific, Cat. No. BP81161), 1 mM of each ribonucleotide (Thermo Fisher Scientific, Cat. No. R0481), 1 U/μl RNase inhibitor (Vazyme, Cat. No. R301-03), 0.001 U/μl pyrophosphatase (Thermo Fisher Scientific, Cat. No. EF0221) and 5% T7 polymerase (home-made). The mixture was incubated at 37 °C for 1 hour, and then an equal amount of polymerase was added to boost transcription for another 30 min. Subsequently, the template DNA was degraded by treatment with 0.14 U/μl Turbo DNase (Invitrogen, Cat. No. AM2238) at 37 °C for 30 min. Finally, the newly transcribed reporter mRNA was isolated using the Monarch RNA Cleanup Kit (New England Biolabs, Cat. No. T2040L), eluted with 1 mM sodium citrate pH 6.4 (Gene Link, Cat. No. 40-5014-05) and quantified as mentioned above.

To generate transcripts containing a 7-methylguanylate cap, we used the Vaccinia Capping System from New England Biolabs (Cat. No. M2080S) according to the manufacturer’s instructions with the addition that the reaction mix was supplemented with 1 U/μl RNase inhibitor (Vazyme, Cat. No. R301-03). The resulting capped mRNA was purified, eluted, and quantified as after the transcription reaction.

### *In vitro* translation

HeLa S3 lysates prepared in hypotonic lysis buffer were used at a concentration of 8.88 ×10^7^ cell equivalents/ml. A standard 12.5 μl reaction was supplemented to a final concentration of 15 mM HEPES, pH 7.3, 0.3 mM MgCl_2_, 24 mM KCl, 28 mM K-acetate, 6 mM creatine phosphate (Roche, Cat. No. 71519-72-7), 102 ng/μl creatine kinase (Roche, Cat. No. 10127566001), 0.4 mM amino acid mixture, unless otherwise stated (Promega, Cat. No. L4461) and 1 U/μl RNase inhibitor (Vazyme, Cat. No. R301-03). To perform *in vitro* translation using lysates prepared in translation-competent buffer, the lysates were directly diluted to 8.88×10^7^ cell equivalents/ml and supplemented with 1 U/μl RNase inhibitor (Vazyme, Cat. No. R301-03). To assess the effect of GADD34 Δ240 and 2-aminopurine (Sigma-Aldrich, Cat. No. A3509) on the phosphorylation status of eIF2α, *in vitro* translation reactions from lysates in translation-competent buffer were supplemented with 0.5 mM GADD34 Δ240 or an equal volume of elution buffer (1x PBS pH 7.5, 250 mM imidazole) dialyzed against protein reconstitution buffer (30 mM NaCl, 5 mM HEPES pH 7.3) and/or 1 to 5 mM 2-aminopurine. Puromycin control reactions contained 320 μg/ml puromycin (Santa Cruz Biotechnology, Cat. No. 58-58-2). Before adding reporter mRNAs, the mixtures were incubated at 33 °C for 5 min, except where otherwise noted. *In vitro* transcribed and capped mRNAs were incubated for 5 min at 65 °C and cooled on ice. Reporter mRNAs were added to the translation reactions at a final concentration of 5 to 80 fmol /μl. The translation reaction was routinely performed at 33 °C for 50 min and extended up to 4 h for time-course experiments and stopped by transferring the samples on ice. RRL *in vitro* translation reactions were carried out according to the manufacturer’s guidelines (Promega, Cat.No. L4960) for 50 min.

### Luciferase Assay

To monitor the protein synthesis output by luciferase assay, a standard 12.5 μl translation reaction was mixed with 50 μl 1x Renilla-Glo substrate in Renilla-Glo assay buffer (Promega, Cat. No. E2720) on a white-bottom 96-well plate (Greiner, Cat. No. 655073). The luminescence signal was measured three times using the TECAN infinite M1000 Pro plate reader according to the manufacturer’s guidelines and plotted on GraphPad (Version 8.4.2). Dual-luciferase assays were performed using 40 μl of the substrate in the appropriate buffer for a 10 μl *in vitro* translation assay (Promega, Cat. No. E1960). Luminescence was measured using TECAN Infinite M1000 and plotted on GraphPad (Version 8.4.2).

### Subcellular fractionation

For the preparation of control cytoplasmic and nuclear lysates, 1×10^7^ HeLa S3 cells were harvested and lysed by resuspension in 1 ml buffer 1 (50 mM HEPES pH 7.3, 140 mM NaCl, 1 mM EDTA, 10% Glycerol, 0.5% NP-40, 0.25% Triton X-100, 1x protease inhibitor cocktail (Bimake, Cat. No. B14002)) and incubation for 10 min at 4 °C rotating. Subsequently, the samples were centrifuged at 500 *g*, 4 °C for 5 min and the supernatant was collected. The same steps were repeated twice with the pellets, and the supernatants were pooled to obtain the cytoplasmic fraction. The pellets were washed by resuspension in 1 ml buffer 2 (10 mM Tris pH 8.0, 200 mM NaCl, 1 mM EDTA, 0.5 mM EGTA, 1x protease inhibitor cocktail (Bimake, Cat. No. B14002)) and incubation for 10 min at 4 °C rotating. The nuclei were pelleted by centrifugation at 500 *g*, 4 °C for 5 min, resuspended in 500 μl buffer 3 (1 mM EDTA, 0.5 mM EGTA, 10 mM Tris pH 8.0, 10% SDS, 0.1% deoxycholate, 1% Triton X-100, 1x protease inhibitor cocktail (Bimake, Cat. No. B14002)) and sonicated with 3x 10 s pulses at 45% amplitude (Vibra-Cell, Cat. No. 75186). Finally, samples were centrifuged at 16,000 *g*, 4 °C for 30 min, and the supernatant was collected to obtain the nuclear fraction.

### Western blot analysis

To assess whether our HeLa S3 lysate is enriched for cytoplasmic factors, samples were collected before lysis by DC (whole cells in hypotonic lysis buffer) and after the final centrifugation step (lysate and pellet). The whole-cell and pellet samples were diluted in three volumes of hypotonic lysis buffer and subjected to sonication for 3x 10 s pulses at 45% amplitude (Vibra-Cell, Cat. No. 75186). Proteins from DC-derived material and the samples deriving from the subcellular fractionation were extracted by chloroform-methanol precipitation. From each sample, 6.8 μg protein (quantification by OD_280_) were loaded.

To compare the phosphorylation status of eIF2α, HeLa S3 cells were harvested from a culture with a density of 0.8×10^6^ to 1×10^6^ cells/ml. Cells were pelleted (200 *g*, 4 °C for 5 min), washed twice with 1x PBS pH 7.4, resuspended in RIPA buffer (50 mM Tris-HCl pH 8.0, 150 mM NaCl, 1% NP-40, 0.5% sodium deoxycholate, 1% SDS) supplemented with protease inhibitor cocktail (Bimake, Cat. No. B14002) and phosphatase inhibitor cocktail (Bimake, Cat. No. B15001), centrifuged at 16,000 *g*, 4 °C for 15 min and the supernatant was collected. HeLa S3 lysates in hypotonic or translation-competent buffer were equally diluted in RIPA buffer. From each sample, approximately 2.5×10^5^ cell equivalents that were adjusted according to the OD_260_ (nucleic acid content in the lysate) were loaded. To assess the phosphorylation status of eIF2α after *in vitro* translation, approximately 3×10^5^ cell equivalents were loaded from each sample. All western blot samples dissolved in 1.5x LDS loading buffer (Invitrogen, NP0008) containing 50 mM DTT were incubated at 75 °C for 5 min. Samples to test nuclear and cytoplasmic markers and eIF2α phosphorylation were run on 4-12% Bis-Tris NuPAGE gels (Invitrogen, Cat. No. WG1401BOX). Samples from *in vitro* translation reactions of hRLuc and 3xFLAG-HBB were run on 4-12% Bis-Tris NuPAGE gels (Invitrogen, Cat. No. WG1203BOX). Samples of SMG6-3xFLAG in vitro translation reactions were run on 3-8% Tris-Acetate NuPAGE gel (Invitrogen, Cat. No EA03752BOX). Transfer onto nitrocellulose membranes was performed using the iBlot 2 Dry Blotting System (Invitrogen, Cat. No. IB21001 and IB23001). The membranes were probed with the following antibodies: Tyrosine-Tubulin (Sigma-Aldrich T9028, 1:10,000), GAPDH (Santa Cruz Biotechnology sc-25778, 1:1,000), Histone H3 (Abcam ab1791, 1:8,000), FLAG M2 (Sigma-Aldrich F3165, 1:2,000), eIF2α (Cell Signaling Technology 9722, 1:1,000), Phospho-eIF2α (Ser51) (Cell signaling Technology 9721, 1:1,000), Renilla luciferase (Thermo Fisher Scientific, Cat. No. PA5-32210, 1:500). To assess proteins with similar molecular weight, the membranes were stripped using 0.2 M NaOH before reprobing.

To quantify the abundance of the synthesized RLuc in the lysates, we established standard curves for quantifying the absolute amounts of RLuc by immunoblotting. Serial dilutions of Recombinant RLuc (RayBiotech, Cat. No. RB-15-0003P) were performed in protein reconstitution buffer (30 mM NaCl, 5 mM HEPES pH 7.3) in the presence of 5% translation competent lysate (to ensure similar sample complexity and electrophoretic resolution to the *in vitro* translation reactions) in 1x LDS buffer (Invitrogen, Cat. No. NP0008). Densitometry was performed using Image Studio Lite Ver 5.2. The average from 3 biological replicates that were run three times (6.67×10^5^ cell equivalents per sample) was calculated by extrapolating the density values of the RLuc standards from the corresponding standard curve that was prepared in Microsoft Excel.

### Production and purification of recombinant GADD34 Δ240

The cDNA encoding the GADD34 protein with a deletion of the amino acids 1-240 was cloned into the pRSET A vector (Thermo Fisher Scientific, Cat. No. V35120) by In-Fusion cloning (Takara, Cat. No. 639650) to generate pRSETA-GADD34 Δ240. The bacterial strain BL-21 (DE-3) (pLysS) was transformed with the cloned plasmids and grown in Luria broth (0.25 l) at 37 °C until the OD600 reached 0.4-0.6. IPTG (1 mM) was then added to induce protein expression, and cells were cultured at 37 °C for 3 h. Bacterial pellets were stored at -20 °C until use. The frozen pellets corresponding to 100 ml bacterial culture were resuspended in 500 μl lysis/wash buffer (1x PBS pH 7.5, 40 mM Imidazole, 1 mM PMSF), lysed by sonication with 3x 10 s pulses at 80% amplitude (Vibra-Cell, Cat. No. 75186) in Eppendorf tubes, and centrifuged at 3,000 *g* for 5 min at 4 °C. The supernatant was mixed with 100 μl HisPur Ni-NTA resin slurry (Thermo Fisher Scientific, Cat. No. 88222) for 3 h at 4 °C under continuous rotation. Unbound proteins were removed by washing the resin 3 times with 600 μl of lysis/wash buffer. The protein was eluted with elution buffer (1x PBS pH 7.5, 250 mM imidazole) and was dialyzed in protein reconstitution buffer (30 mM NaCl, 5 mM HEPES pH 7.3) overnight using a Slide-A-Lyzer Dialysis Cassette (MWCO 3.5K) (Thermo Fisher Scientific, Cat. No. 88400TS).

## RESULTS AND DISCUSSION

### Dual centrifugation as a method to produce translation-competent lysates

The efficient lysis of eukaryotic cells under well-defined biochemical conditions is a prerequisite for a wide range of biological assays. The lysis process is often challenging, especially when the isolation of active biomolecules is required for downstream applications. While many different protocols have been developed to produce biologically active lysates that can be used for *in vitro* studies, they are often limited in accurately defining the mechanical force applied during lysis. To apply reproducible shearing forces, we opted for dual centrifugation (DC). In this process, the samples undergo a simultaneous rotation around two axes of symmetry (Fig. 1A). The continuous change of centrifugal forces in the sample vials leads to shearing and homogenization of the processed material (17,18). We rationalized that such a mechanical treatment would allow the lysis of cells independently of the buffer conditions. Importantly, this method accurately defines and fine-tunes the lysis parameters by controlling the centrifugation speed while keeping the processed material at a low temperature.

Based on our previously established *in vitro* translation assays (16), we produced translation-competent lysates using DC as a cell lysis method. The steps of our protocol, as illustrated in Fig. 1B, are the following: **1**. Dual centrifugation of cells in hypotonic lysis buffer **2**. Centrifugation to remove cell debris and nuclei **3**. Retention of the supernatant (lysate), which can be stored frozen for further use **4**. Supplementation of the lysate with a buffer mixture containing salts, energy sources and amino acids required for translation **5**. Addition of reporter mRNAs **6**. *In vitro* translation followed by detection of the protein product or other downstream approaches.

### Dual centrifugation can lyse human cells and allows for the isolation of cytoplasm-enriched extracts

Dual centrifugation has not been previously used to disrupt living cells and isolate biologically active lysates. We set out to assess whether cell lysis can, in principle, be performed using this method. We established our system using HeLa S3 cells (spinner cells), which are adapted to grow in suspension culture and reach high cell densities. The cells were harvested at a density of 0.8 – 1.0×10^6^ cells/ml in suspension, resuspended in a hypotonic lysis buffer to reach a final concentration of 2×10^8^ cells/ml and subjected to DC. To allow a comparative analysis, the dual centrifugation experiments were performed for 4 min and the temperature of the rotor was set at -5 °C, taking into consideration a possible increase of the sample temperature during DC. In all experiments, it was confirmed that the samples did not freeze during this step.

First, we assessed under which centrifugation speed cell lysis can be achieved, ideally without disrupting the cell nuclei. To visualize the lysis efficiency, we performed a Trypan Blue staining that assesses cell membrane integrity; living cells are visible as white spheres, whereas dead cells and nuclei as dark blue foci. As shown in Fig. 2A, the lysis efficiency increased proportionally to the applied centrifugal force. Lysis was achieved even under isotonic buffer conditions (in PBS, data not shown), rendering dual centrifugation an appealing method for lysing cells in virtually any buffer conditions without the addition of detergents. This is of particular interest in cases where preserving intact, functional biomolecules is crucial. Since we wanted to obtain translation-competent lysates, we opted for a mainly cytoplasmic extract and applied dual centrifugation forces in the range of 250 – 1000 RPM to titrate the translation efficiency of the resulting lysates.

**Figure 2:**
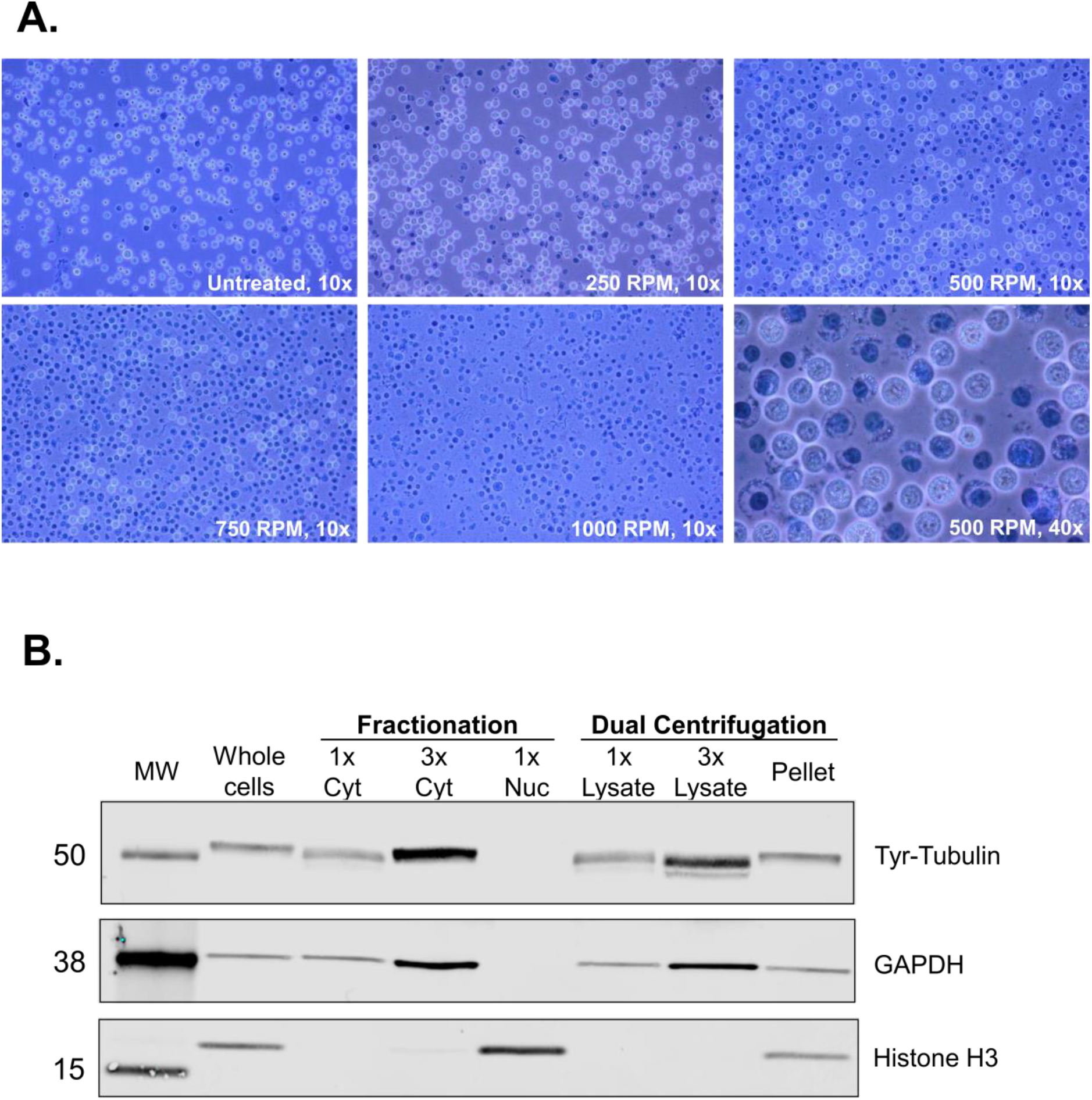
Dual centrifugation allows for the production of cytoplasm-enriched lysates. **A**. Assessment of lysis efficiency by Trypan Blue staining under the light microscope at 10x or 40x magnification. **B**. Western blot analysis of cytoplasmic and nuclear markers of the supernatant (lysate) and the pellet of DC-treated cells. Cytoplasmic and nuclear fractions were prepared using an alternative method of biochemical fractionation. All samples correspond to 6.8 μg protein.

After DC, we removed residual intact cells, nuclei and insoluble fragments from the lysate by regular centrifugation. To assess whether our lysates are enriched for cytoplasmic components, we assessed cytoplasmic and nuclear markers by immunostaining. We compared our lysates (supernatant after DC using 500 RPM and regular centrifugation, as shown in Fig. 1B) with total cell lysates and cytoplasmic and nuclear lysates obtained using a detergent-based protocol for subcellular fractionation. As shown in Fig. 2B, the supernatant is enriched for cytoplasmic markers (Tyrosine-Tubulin and GAPDH). No signal for histones could be detected even when loading the triple amount of supernatant. The pellet contains both nuclei and cytoplasmic components due to the incomplete cell lysis under mild conditions. These results show that dual centrifugation allows for the production of lysates that are highly enriched for cytoplasmic components. Further optimization could allow for the application of this method to separate cytoplasmic from nuclear fractions.

### Optimization of the conditions for *in vitro* translation using dual centrifugation-derived lysates

To assess whether the DC-derived lysates can translate reporter mRNAs, we used an *in vitro* transcribed and capped *Renilla* luciferase reporter that contains an 80-nucleotide long poly(A) tail (RLuc reporter). To this end, we set up *in vitro* translation reactions for lysates derived from DC at 250, 500, 750, or 1000 RPM. The processed lysates (i.e., the supernatant after the final, regular centrifugation) were supplemented with a buffer containing necessary salts, energy source and amino acids needed for *in vitro* translation and with an equal amount of mRNA for the four reactions. After a 50 min incubation at 33 °C, we measured the enzymatic activity of RLuc, which is proportional to the protein production in the lysates. As shown in Fig. 3A, the luminescence measurements verify the translation capacity of all the tested lysates, with an optimal activity of lysates that were subjected to DC at 500 RPM and underwent, as shown in Fig. 2A, a relatively mild lysis process. No translation activity was detected in the presence of the translation inhibitor puromycin. The fact that milder lysis conditions lead to a higher translation output may be attributed to the optimal balance between disruption of sensitive components due to the mechanical forces and the efficiency of releasing the cytoplasm. Additionally, the prevention of lysis of nuclei may also contribute to the higher translation efficiency of the lysates.

**Figure 3:**
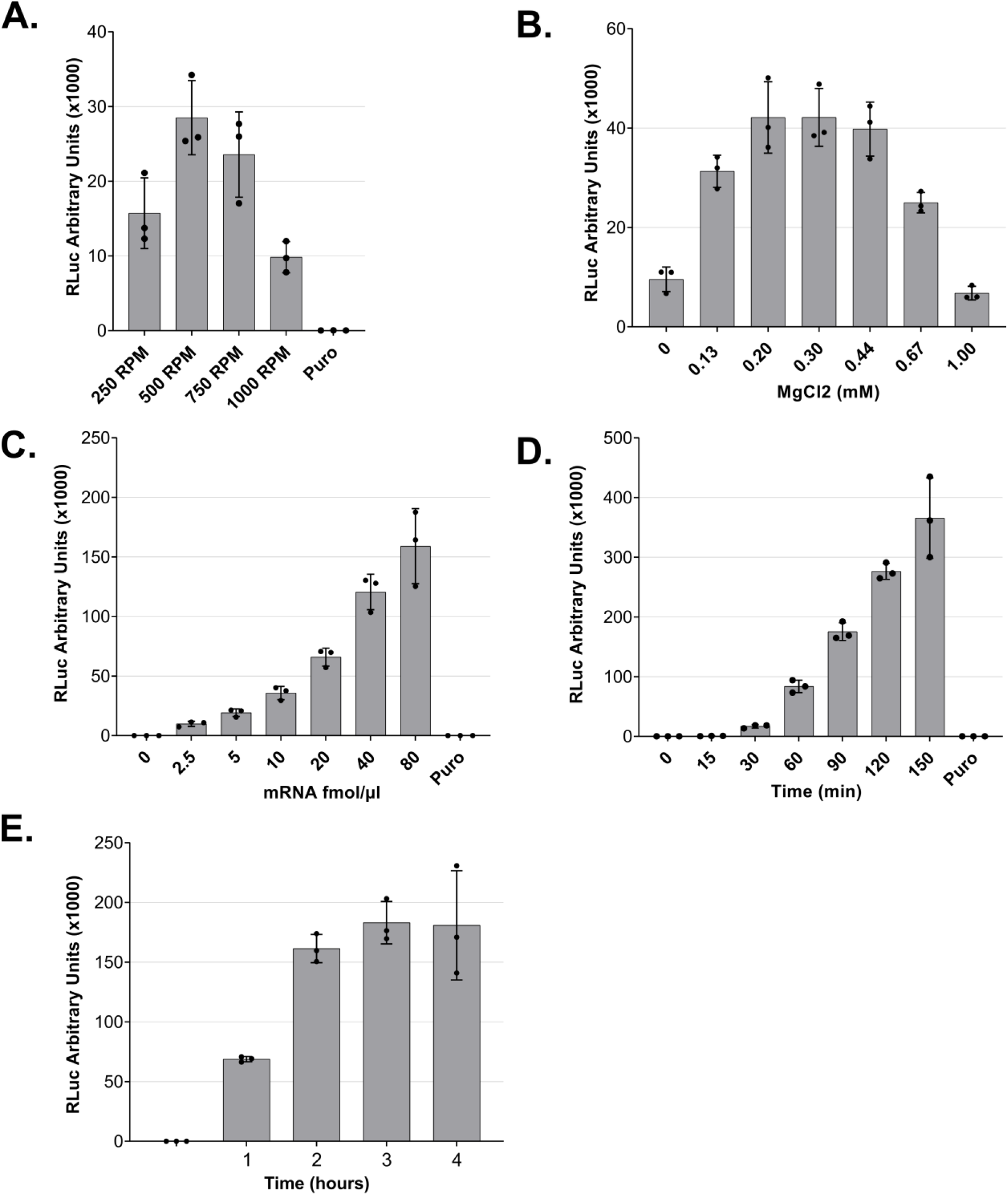
Optimization of lysate preparation and *in vitro* translation. **A**. *In vitro* translation using lysates that were produced under different DC conditions. **B**. Mg^2+^ titration for *in vitro* translation reactions. **C**. *In vitro* translation using different mRNA concentrations. **D and E**. Time-course of RLuc *in vitro* translation reactions. For all experiments (except C), 5 fmol/μl of RLuc mRNA was used. For all panels, RLuc activity measurements are depicted as arbitrary units (AU) of luminescence. Each dot depicts the value of an individual experiment, and *in vitro* translation (biological replicate) was measured three times (technical replicates). Mean values and SD are shown.

To test our DC-derived lysates, we applied the *in vitro* translation conditions as they were previously optimized for our syringe-treated lysates (16). Among the most crucial factors for efficient translation is the Mg^2+^ concentration. Thus, we decided to titrate this parameter to ensure that the concentration is optimal under the new lysis conditions. Setting as reference a concentration of 0.30 mM, we assessed the efficiency of translation under different MgCl_2_ concentrations in the translation buffer, and we observed that the concentration of choice falls in the optimal range of translation activity (Fig. 3B). For this reason, we selected a concentration of 0.30 mM MgCl_2_ for our *in vitro* experiments. Notably, the range of suitable concentration was wider than for our previous translation system, indicating that lysates prepared by DC are more robust towards small changes of *in vitro* translation conditions.

Next, we titrated the effect of the reporter mRNA concentration on the translation output of the *in vitro* translation reaction. We added increasing amounts of reporter mRNA, ranging from 0 to 80 fmol/μl of reaction, and as shown in Fig. 3C, we observed a concentration-dependent increase of the reporter protein output. This increase is linear for concentrations up to 40 fmol/μl (Fig. 3C). Based on this result, we routinely use 5 or 10 fmol/μl for standard translation reactions since these concentrations clearly correspond to the linear range and therefore do not saturate the translation system with mRNA. The concentration of mRNA used can be adjusted depending on the downstream applications and depends on the translation efficiency of the individual mRNA.

To assess how the duration of the incubation affects the production of the recombinant protein and, therefore, the translation efficiency of the system, we performed a time course from 0 to 150 min (Fig. 3D) and 0 to 4 h (Fig. 3E). We observed an almost linear increase of the luminescence signal during the entire 150 min incubation. When we prolonged the assay duration, the reporter protein production reached a plateau after 2 h of translation. Interestingly, there is a non-linear increase of the signal during the first 30 min of incubation, in accordance with the observations that the initial association of translation factors with an mRNA is a rate-limiting step in translation (21).

### Dual centrifugation-derived lysates can be used for cap- and IRES-dependent *in vitro* translation

Cap-dependent translation is the primary mechanism of translation initiation for mammalian mRNAs. For this reason, we decided to address the effect of a 5’ cap on the translation efficiency of mRNA reporters in the *in vitro* translation system by comparing the RLuc output of *in vitro* capped and uncapped *Renilla* luciferase reporter mRNAs. We used equimolar amounts of the two mRNA molecules and as shown in Fig. 4A, we observed a >30-fold increase in the efficiency of translation in the presence of a 5’ cap on the reporter mRNAs. This result shows that our system is sensitive to the presence of a cap, in accordance with the cap-dependent translation initiation of most physiological mRNAs in mammalian cells.

**Figure 4:**
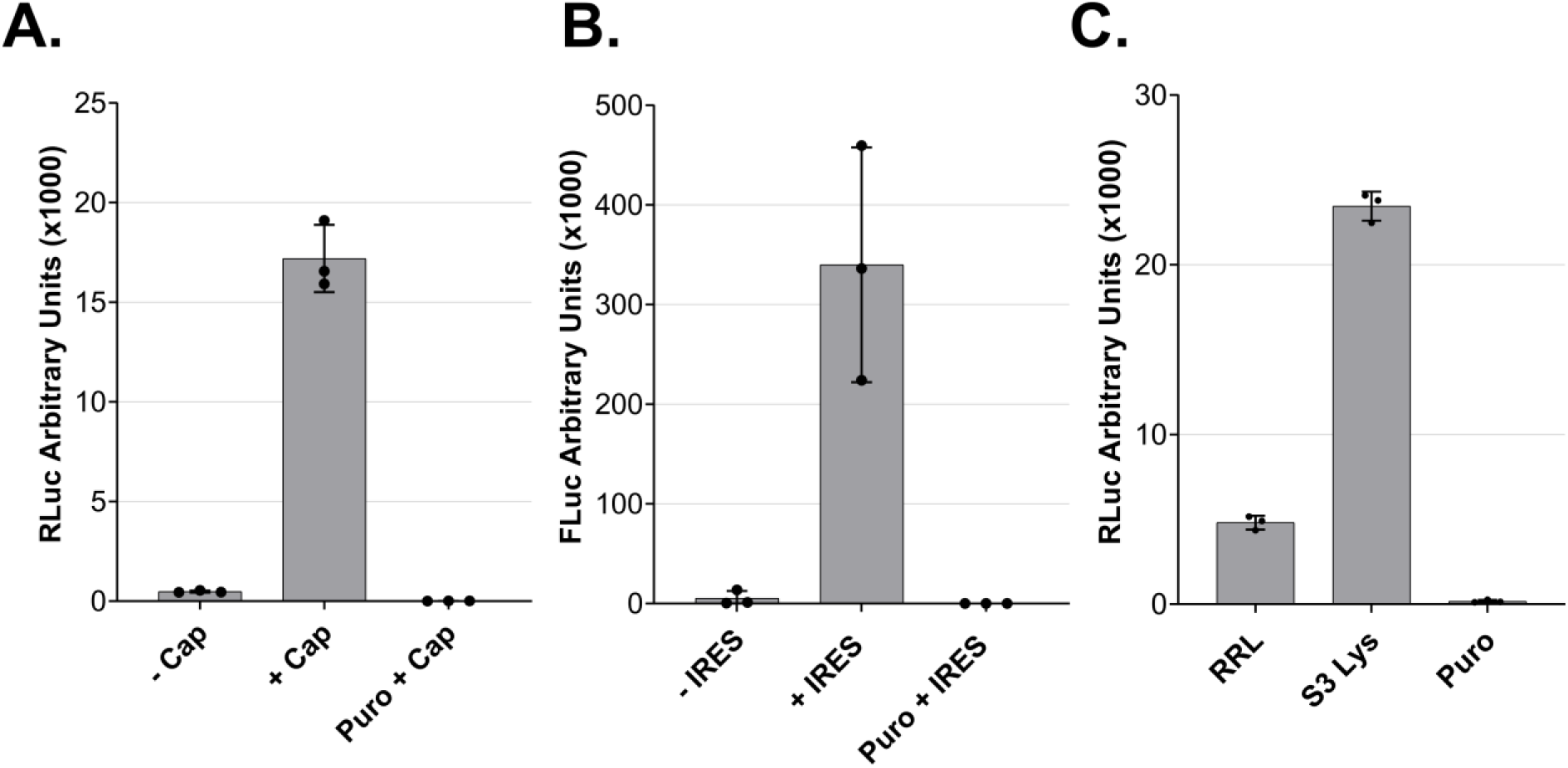
Cap- and IRES-dependent translation of dual centrifugation-derived lysates and comparison to rabbit reticulocyte lysate. **A**. Comparison RLuc measurements of *in vitro* translation reactions of DC-generated lysates with uncapped (-Cap) or capped (+Cap) mRNAs. **B**. Comparison of Firefly Luc (FLuc) measurements of *in vitro* translation reactions of DC-generated lysates using dual-luciferase reporters with or without an ECMV-IRES upstream of the FLuc ORF. **C**. *In vitro* translation using Rabbit reticulocyte lysate (RRL) and three HeLa S3 lysates. Luc activity measurements are depicted as arbitrary luminescence units (AU). The dots depict the values of three individual experiments (biological replicates) that were measured each three times (technical replicates). *In vitro* translation experiments were performed using 5 fmol/μl mRNA reporters.

After optimizing the translation efficiency of cap-dependent translation, we addressed whether DC-derived lysates can perform IRES-dependent translation, a cap-independent process (22). IRES usage is of interest because many vectors have been developed for *in vitro* protein production employing this principle. To address whether we can perform IRES-dependent translation in our lysates, we prepared a dual-luciferase reporter mRNA encoding *Renilla* and Firefly Luciferase. We used two versions of this construct, one where the two ORFs are separated by an IRES originating from the human EMCV virus and a control version that lacks the IRES sequence (23). The detection of a clear Firefly luciferase signal only in the presence of the IRES demonstrates the capacity of the lysate to perform IRES-dependent translation (Fig. 4B).

We compared the efficiency of our system to a commercially available rabbit reticulocyte lysate system (RRL). The *in vitro* translation reaction in the RRL was performed according to the manufacturer’s guidelines with an equal amount of RLuc reporter. As shown in Fig. 4C, our lysate is about five-fold more translationally active than the RRL. Thus, our system allows for the reproducible production of translation-competent lysates with high translation potential.

### Direct cell lysis in a translation-competent buffer

The lysate described above was prepared using a hypotonic lysis buffer complemented with the necessary salts, amino acids and energy regeneration system before *in vitro* translation. We decided to assess whether lysates can also be prepared in a buffer that already contains all the necessary components. When we lysed the cells directly in such a translation-competent buffer, we obtained translationally active lysates with a similar activity compared to the previous method of lysate preparation (Fig. 5A). Moreover, we observed that the translation is enhanced if it is not preceded by a 5 min pre-incubation at 33 °C before adding the reporter mRNA under these conditions (Sup. Fig. 1).

**Fig. 5:**
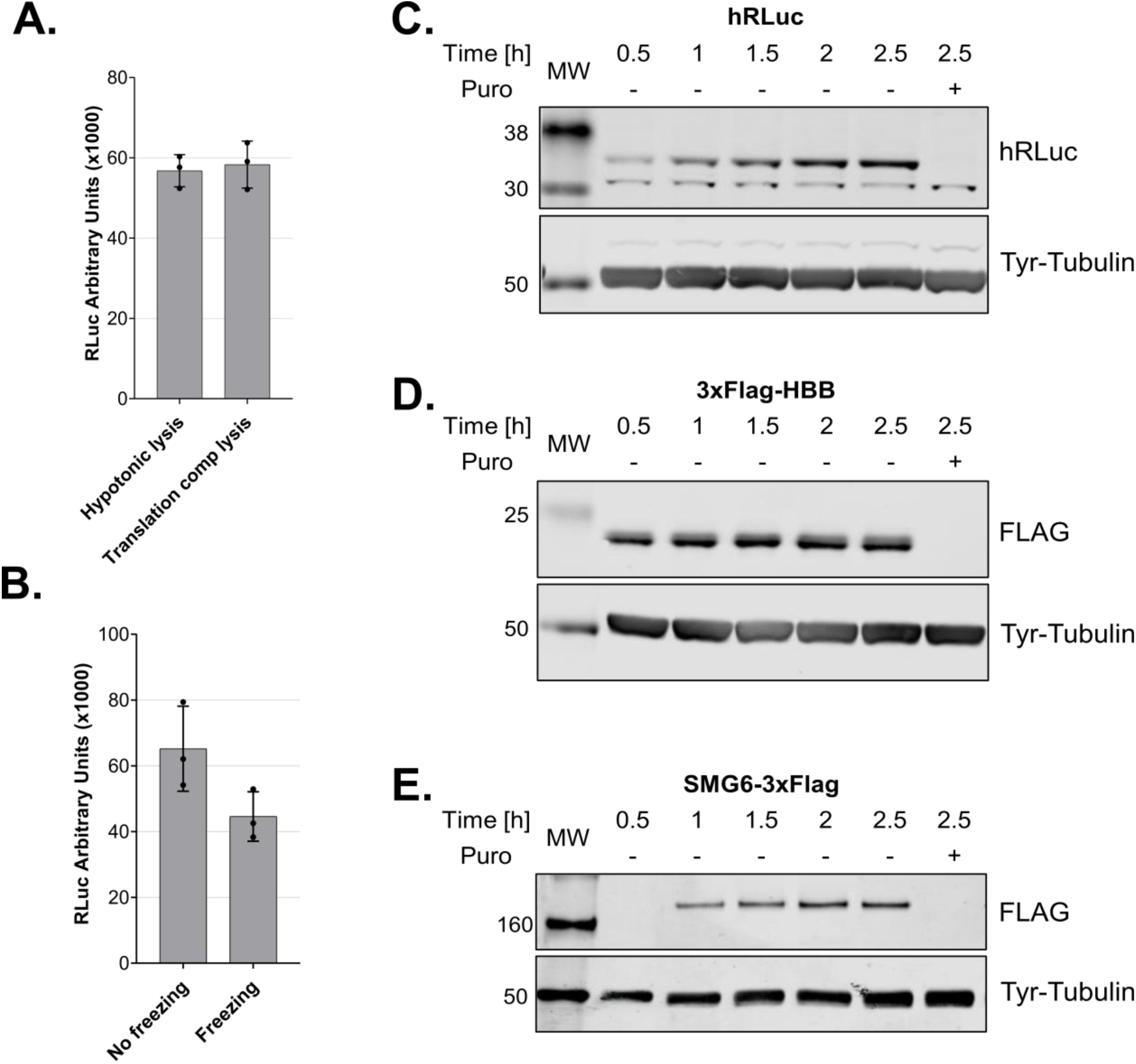
Cell lysis in a translation-competent buffer and of frozen cells and *in vitro* translation of different mRNA reporters. **A**. Comparison of lysates produced using a hypotonic lysis buffer complemented with components required for *in vitro* translation (hypotonic lysis) versus lysis in a translation-competent buffer (translation comp lysis). **B**. Comparison of lysates produced using a hypotonic lysis buffer complemented with components required for *in vitro* translation (hypotonic lysis) from freshly harvested cells versus from frozen cell pellets. For A and B, RLuc activity measurements are depicted as arbitrary luminescence units (AU). Each dot depicts the value of an individual experiment and *in vitro* translation (biological replicate) that was measured three times (technical replicates). Mean values and SD are shown. **C-E**. Western blot analysis of time-course of *in vitro* translations of different mRNA reporters using lysates produced in translation-competent buffer. For C and D 1.6 μl and for E 1.25 μl of the translation reactions were loaded on the gel. *In vitro* translation experiments were performed using 10 fmol/μl RLuc reporter, a final amino acids concentration of 0.01 mM and without preincubating the translation reactions before adding the reporter mRNAs.

Additionally, we assessed whether it is possible to generate active lysates using pellets from frozen cells. We compared lysates from the same amount of frozen or fresh cells prepared in a translation-competent buffer. The lysates deriving from frozen cells were translationally active, although with reduced activity compared to unfrozen cells (Fig. 5B).

The cell lysis in a translation-competent buffer further facilitates the production of translation-competent lysate and improves the reproducibility of the *in vitro* translation assay by reducing intermediate steps. Such an approach is ideal for structural studies that require large amounts of translationally active lysates. The fact that we can obtain translationally active lysates from frozen cells allows us to decouple cell culturing and harvesting from the lysate production, which further expands the range of possible applications. Under these conditions, we also assessed whether it is possible to translate reporter mRNAs with different 5’UTRs and ORFs encoding proteins with different molecular masses. We cloned the ORF of human beta globin (HBB) and SMG6, an endonuclease that is implicated in nonsense-mediated mRNA decay (24–26), and used the corresponding *in vitro* transcribed capped and polyadenylated mRNAs, which harbor the SARS-CoV-2 leader sequence in the 5’UTR, as templates for *in vitro* translation. Similarly to RLuc (36 kDa, Fig. 5C), we observed single distinct bands for 3xFlag-HBB (21 kDa, Fig. 5D) and SMG6-3xFlag (163 kDa, Fig. 5D).

### eIF2α dephosphorylation in the lysates increases the translation output

To use our system for the production of recombinant proteins, maximizing the translation efficiency is crucial. The eukaryotic initiation factor 2 (eIF2) is a key regulator of global translation initiation (21) and the dephosphorylation of its α subunit (eIF2α) has been shown to increase the translation efficiency of HeLa lysates (27). We assessed and modulated the phosphorylation status of eIF2α in our lysates to further increase the protein production capacity. eIF2 is the major Met-tRNAi carrier when it is bound to GTP, however, eIF2 is more stable when it is bound to GDP (28). For this reason, the activation of eIF2 requires the activity of a guanine exchange factor, eIF2B, which promotes GDP dissociation and facilitates GTP and Met-tRNAi to form a ternary complex with eIF2. eIF2 is subject to tight control as it can be phosphorylated by several protein kinases at Ser 51 of its α subunit (eIF2α). Once phosphorylated, eIF2α does not allow eIF2B to exchange GDP to GTP, and protein synthesis initiation is inhibited (21).

To assess the phosphorylation status of eIF2α in our cells and lysate, we immunoblotted total and Ser51-phosphorylated eIF2α. We observed that eIF2α was considerably phosphorylated in the lysates compared to intact cells before lysis (Fig. 6A), probably due to the activity of kinases during and/or after lysis. To modulate the phosphorylation status of eIF2α in the lysates during *in vitro* translation, we used GADD34 and 2-aminopurine. GADD34 forms a complex with the catalytic subunit of protein phosphatase 1 (PP1) and promotes the dephosphorylation of eIF2α *in vivo* (29). It was shown that treatment with a 240 amino acids N-terminal truncation of recombinant GADD34 (GADD34 Δ240) can reduce the phosphorylation state of eIF2α and increase translation efficiency of HeLa lysates (27). 2-aminopurine (2-AP) is an inhibitor of the protein kinase R (PKR) and other serine/threonine protein kinases and therefore represses the phosphorylation of eIF2α at Ser51 (30,31).

**Figure 6:**
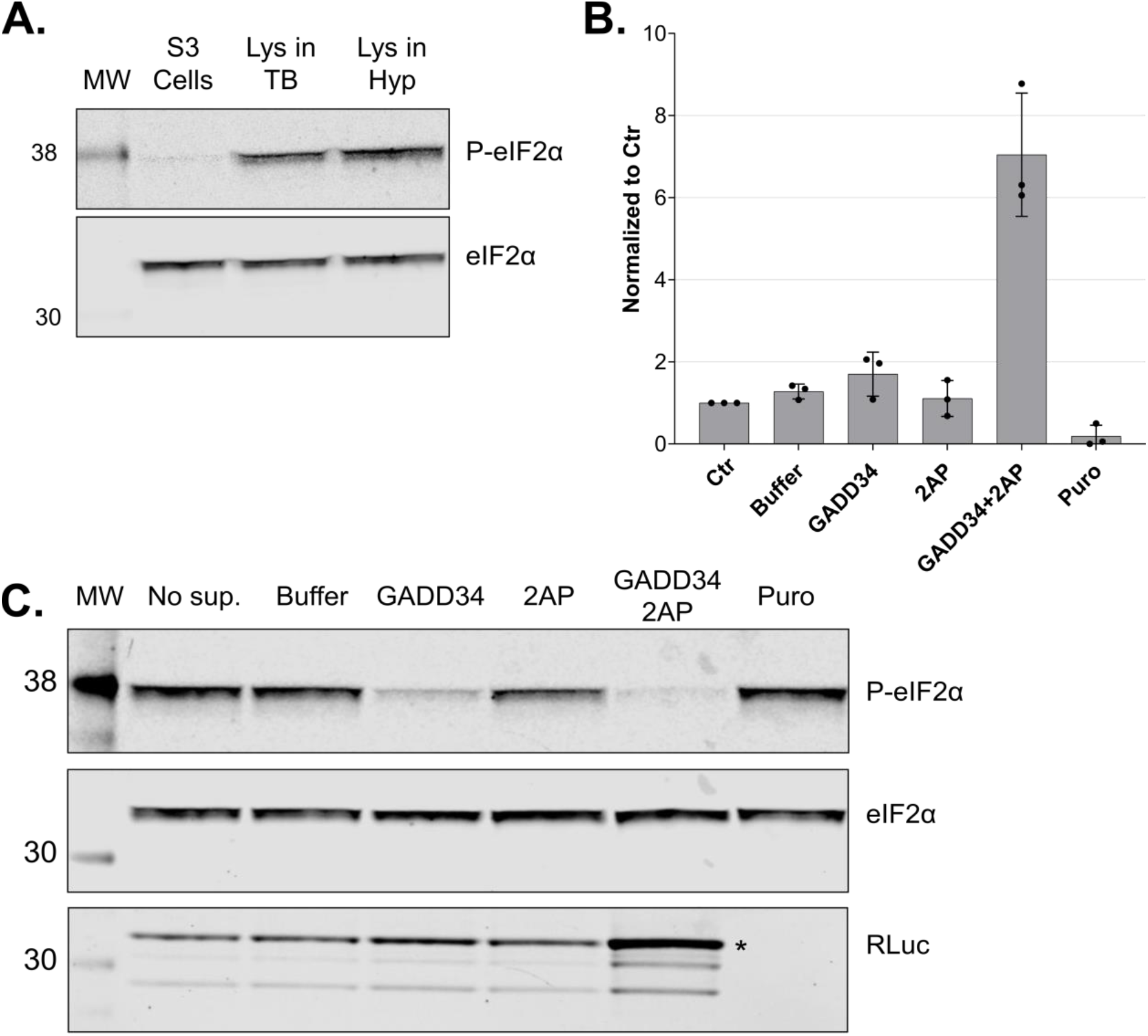
*In vitro* dephosphorylation of eIF2α leads to increased translation output. **A**. Western blot analysis of extracts from growing cells and translation-competent lysates against eIF2α and P-eIF2α (Ser51). All samples correspond to approx. 2.5×10^5^ cell equivalents. **B**. Comparison of RLuc measurements of *in vitro* translation reactions of DC-derived lysates in the absence (Control) or the presence of 0.5 mM GADD34 Δ240 and 3 mM 2-aminopurine. RLuc activity measurements are depicted as arbitrary units (AU) of luminescence. The dots depict the values of three individual experiments (biological replicates) that were measured each three times (technical replicates). **C**. Western blot analysis of lysates after *in vitro* translation against eIF2α and P-eIF2α of non-supplemented lysate (No sup.), the addition of protein reconstitution buffer (Buffer), 0.5 mM GADD34 Δ240, 3 mM 2-aminopurine, 0.5 mM GADD34 Δ240 in combination with 3 mM 2-aminopurine or treated with puromycin. All samples correspond to 3×10^5^ cell equivalents. *In vitro* translation experiments were performed in lysates produced using translation-competent buffer using 10 fmol/μl RLuc reporter without preincubating the translation reactions before adding the reporter mRNAs. Asterisk denotes the position of full-length RLuc.

To reduce the phosphorylation of eIF2α in our lysates, we expressed and purified recombinant GADD34 Δ240 (Sup. Fig. 2A), titrated 2-AP in the presence of the optimal GADD34 Δ240 concentration, and assessed the effect on translation using RLuc reporter assays (Sup. Fig. 2B). The addition of 2-AP alone led to a decrease in eIF2α phosphorylation but did not affect the translation output. The addition of GADD34 Δ240 led to a distinct decrease in eIF2α phosphorylation and a two-fold increase in the translation output (Figure 6B and C). Interestingly, the addition of both 2-AP and GADD34 Δ240 (3 mM and 0.5 mM, respectively) to the translation reactions led to almost complete dephosphorylation of eIF2α (Fig. 6B) and a 6-fold increase of protein synthesis compared to the untreated lysates (Fig. 6C).

We quantified the protein output of RLuc under these conditions by comparing the immunoblot signal of the produced protein to standard dilutions of recombinant RLuc and estimated the production of approximately 1 ng of recombinant protein per μl of lysate after 50 min of translation (Sup. Fig. 2C). This output can be further increased by prolonging the translation reaction, increasing the concentration of the reporter mRNA, and using more concentrated lysates of higher cell equivalent density (data not shown). In other words, the translation-competent lysates can be used to produce potentially toxic proteins *in vitro*, yielding more than 1 μg protein per milliliter of translation reaction.

In conclusion, we provide a new protocol for producing translation-competent lysates from human cells in a fast, scalable, and highly reproducible manner. Our system is highly active and enables cap-dependent translation with unprecedented protein yields, which not only facilitates fundamental research on translation regulation but also paves the way for the future development of large-scale *in vitro* protein synthesis in a human system.

## ACKNOWLEDGEMENTS

We thank Prof. Ulrich Massing and his team at Hettich AG for fruitful discussions about the possible applications of DC and their introduction to the ZentriMix 380 R system. We also thank Andrea Eberle for her technical advice on the project and the critical reading of the manuscript, Nicole Kleinschmidt for her technical advice, and Sofia Nasif for her remarks on the manuscript.

## SUPPLEMENTARY MATERIAL

**Sup. Fig. 1:**
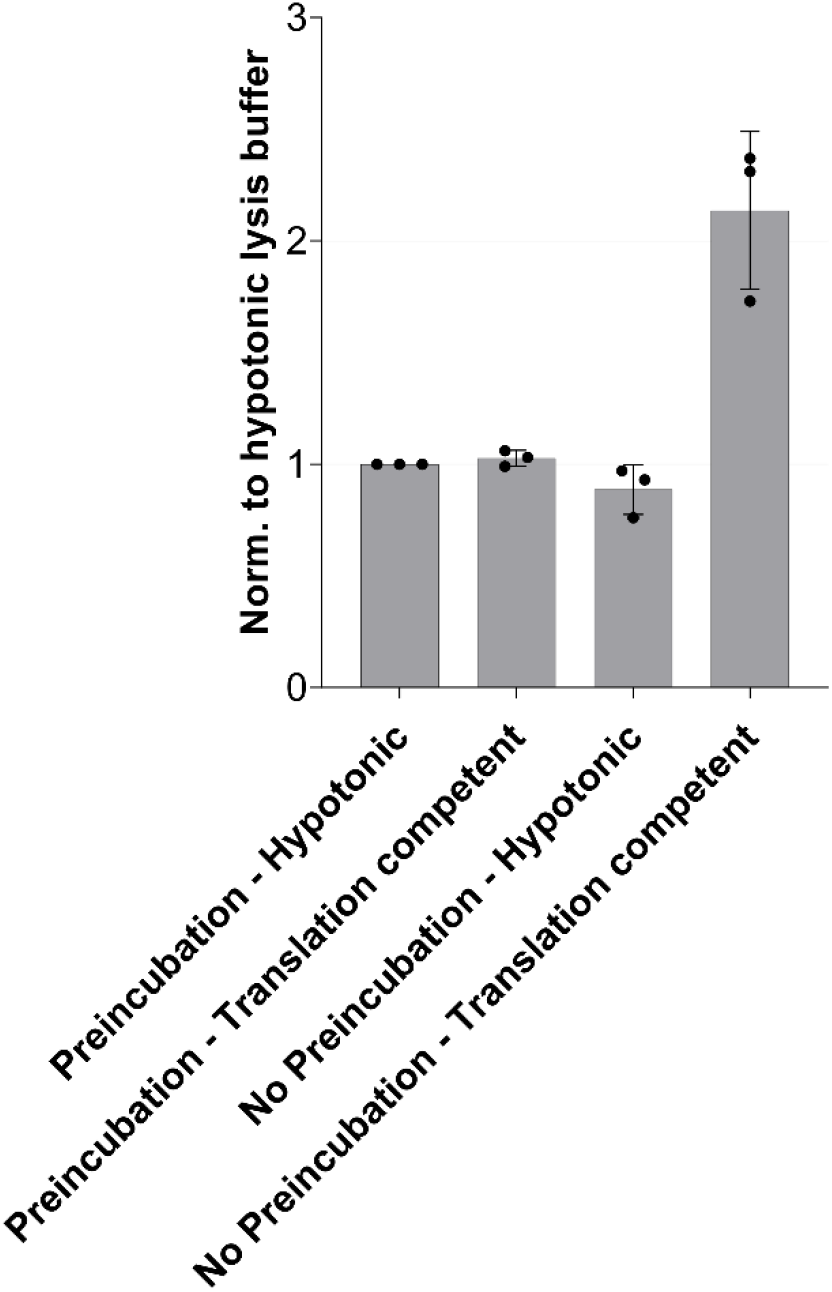
Comparison of lysates produced using a hypotonic lysis buffer complemented with components required for *in vitro* translation (hypotonic) versus lysis in a translation-competent buffer (translation comp lysis) with or without a 5’ pre-incubation at 33 °C before *in vitro* translation. RLuc activity measurements are depicted as normalized values of luminescence to the hypotonic buffer condition with pre-incubation. Each dot depicts the value of an individual experiment and *in vitro* translation (biological replicate) that was measured three times (technical replicates). *In vitro* translation experiments were performed using 10 fmol/μl RLuc reporter. Mean values and SD are shown.

**Sup. Fig. 2:**
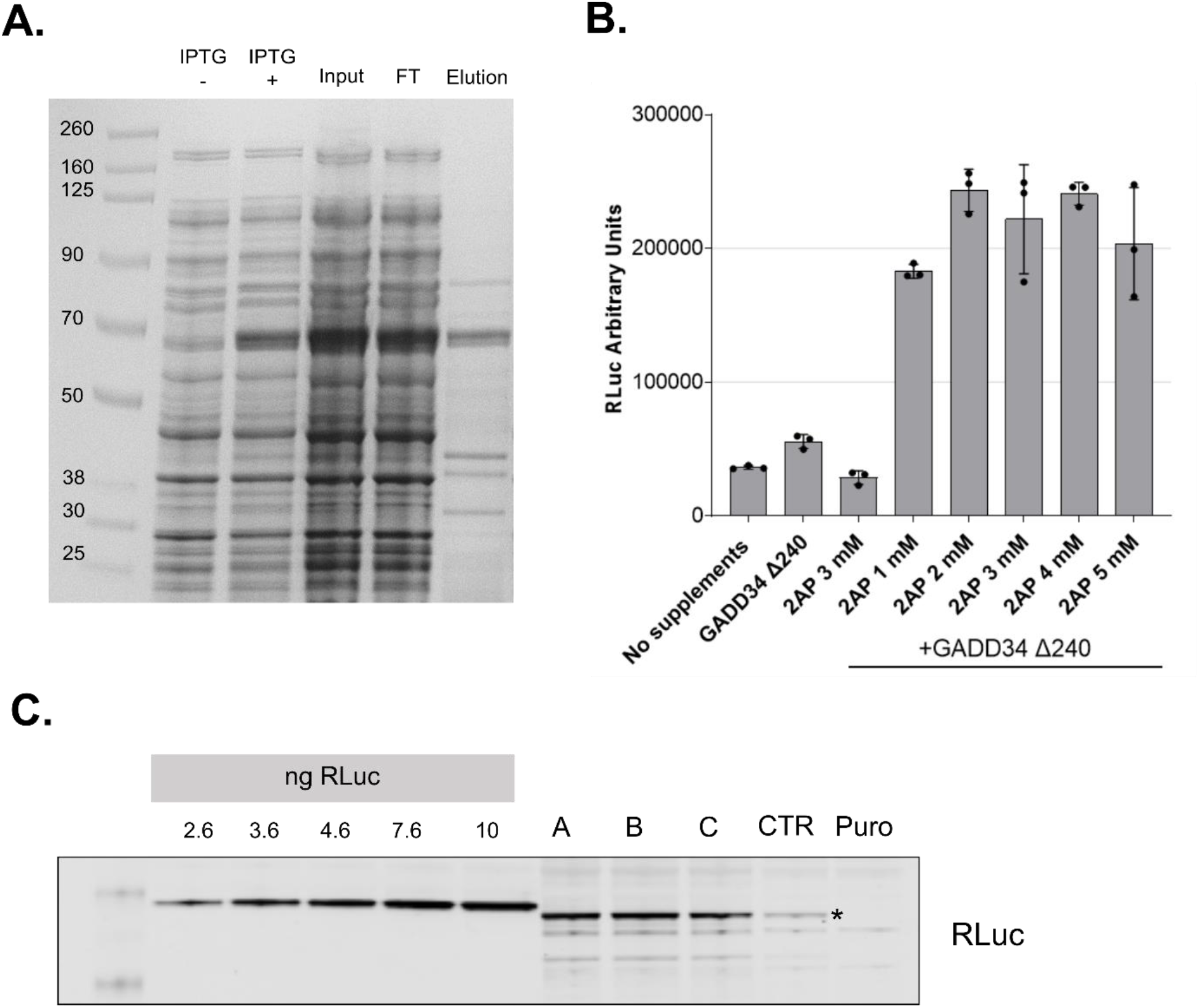
**A**. Coomassie-stained gel showing the purification steps of GADD34 Δ240. Bacterial extract in the presence or absence of IPTG in the culture. Samples (5 μl) of input, flowthrough and purified GADD34-His fractions were analyzed. **B**. Comparison of RLuc measurements of *in vitro* translation reactions of DC-treated lysates in the absence (No supplements) or the presence of 0.5 mM GADD34 Δ240, 3 mM 2-aminopurine and increasing concentrations of 2-aminopurine in the presence of 0.5 mM GADD34 Δ240. RLuc activity measurements are depicted as arbitrary units (AU) of luminescence. The dots depict the values of three individual experiments (biological replicates) that were measured each three times (technical replicates). **C**. Western blot analysis of increasing amounts of recombinant RLuc diluted in translation competent and lysates corresponding to 6.67×10^5^ cells after *in vitro* translation against RLuc of non supplemented lysate (CTR), or three replicates (A, B, and C) of lysates supplemented with 0.5 mM GADD34 Δ240 and 3 mM 2-aminopurine. *In vitro* translation experiments were performed using 10 fmol/μl RLuc reporter without preincubating the translation reactions before adding the reporter mRNAs.

**Sup. Fig. 3.**
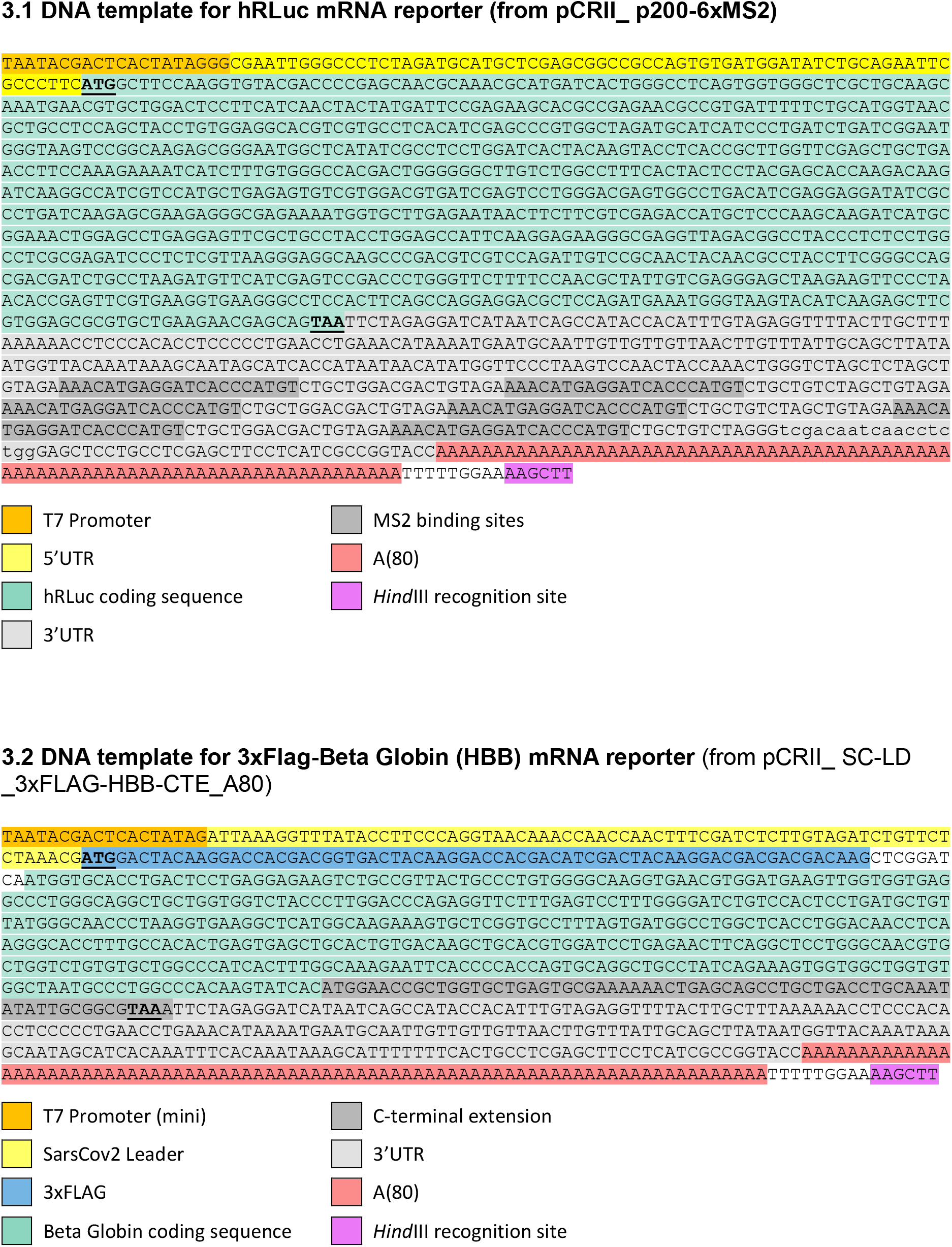
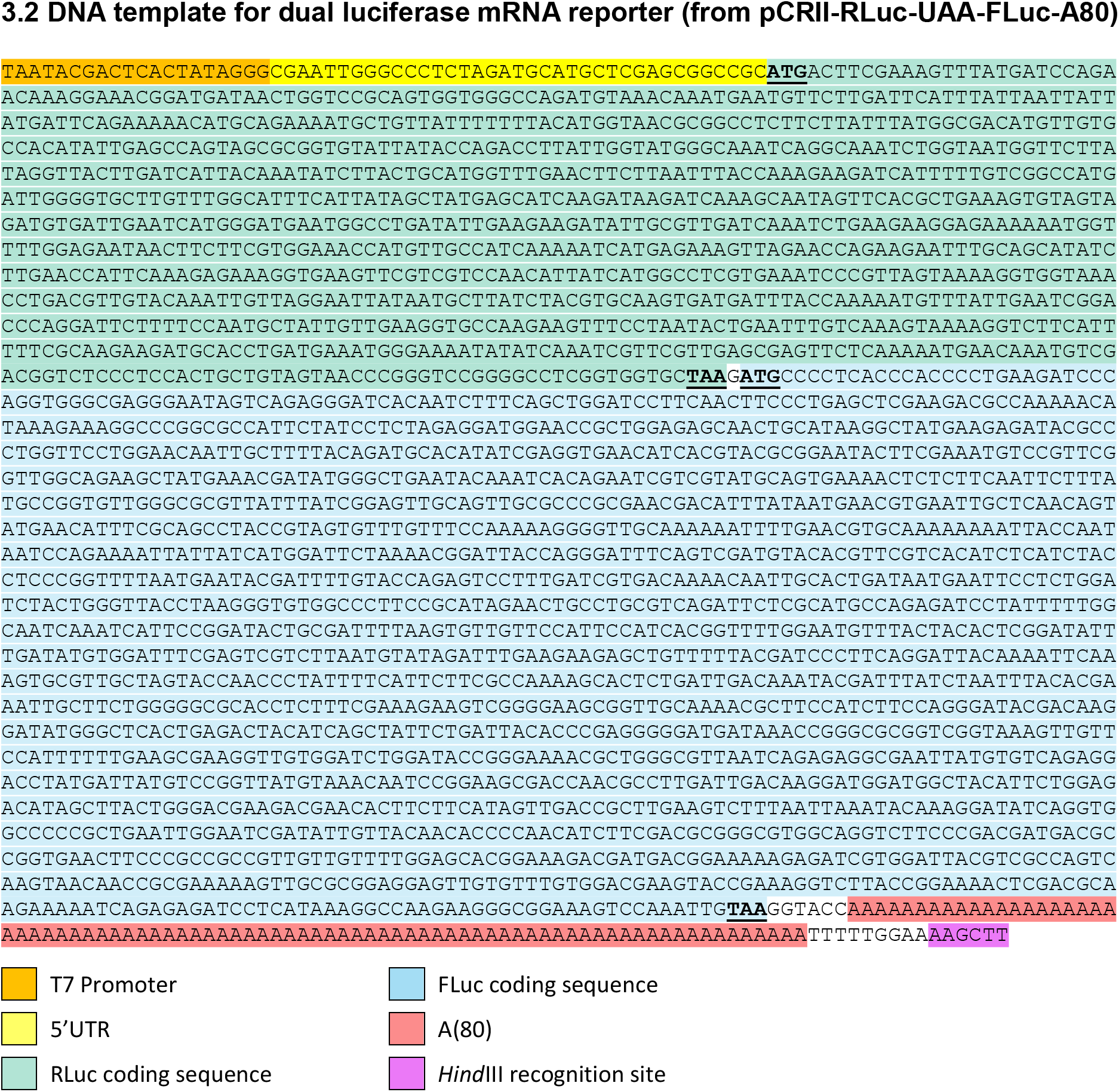
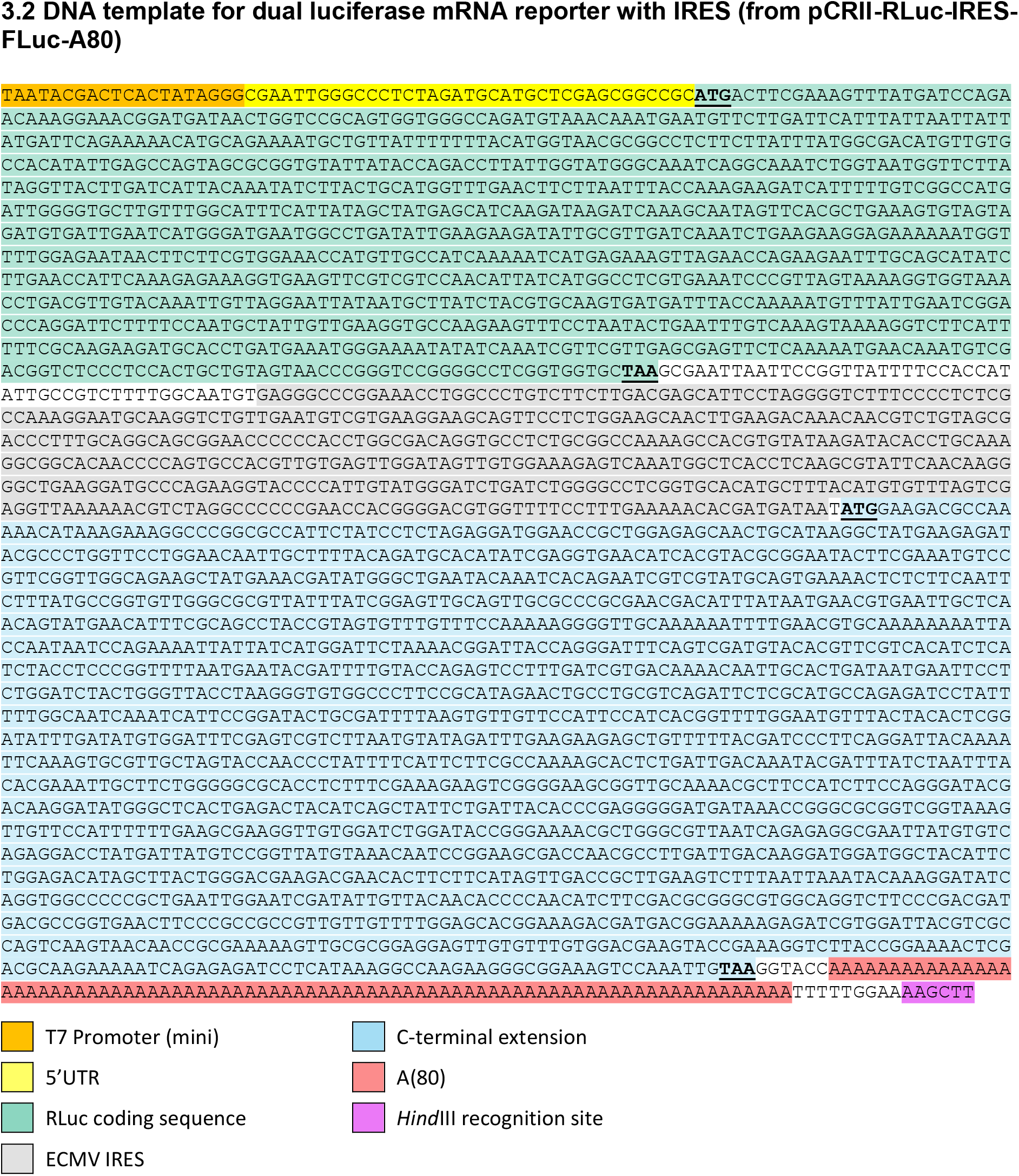
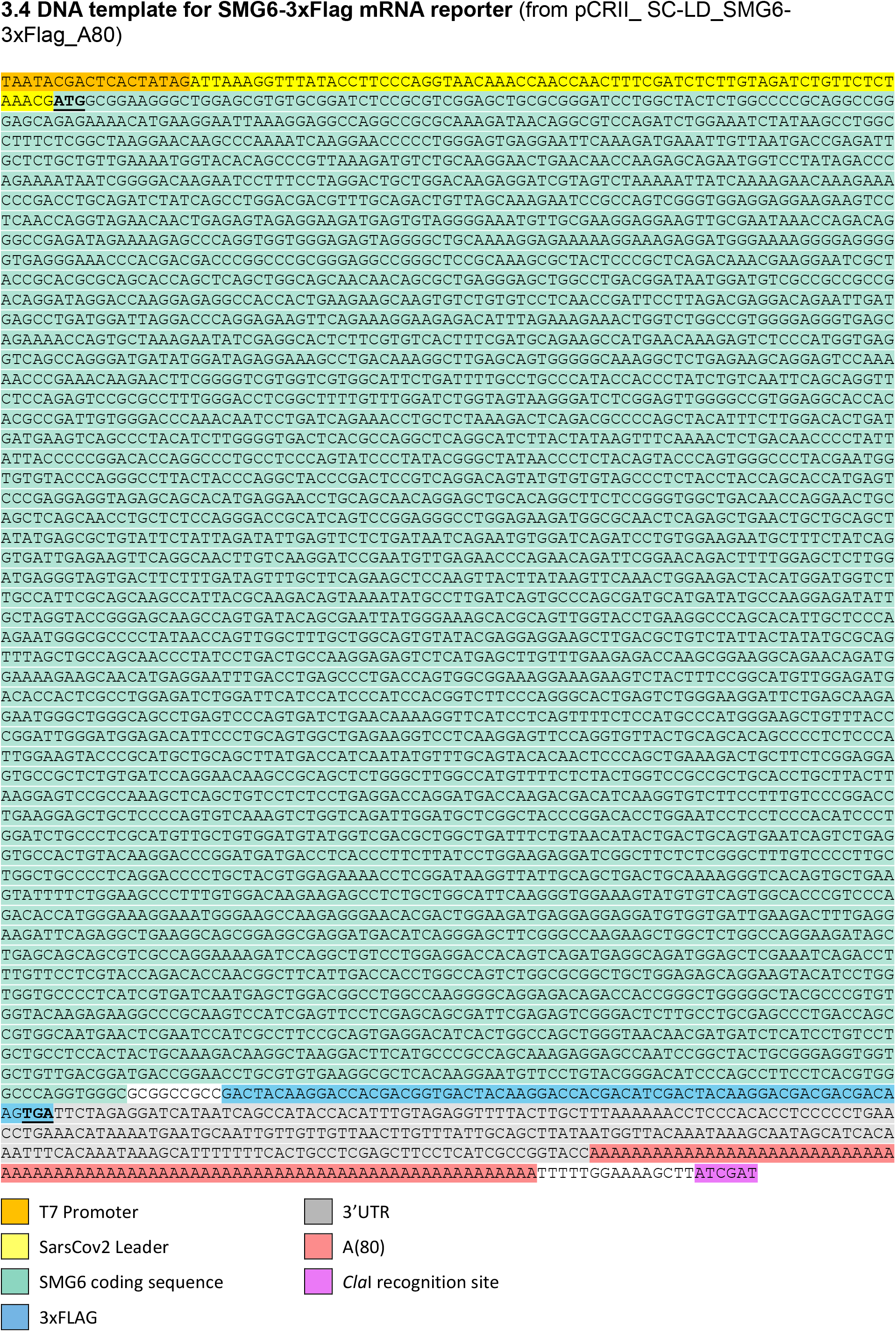
Nucleotide sequences of the DNA templates used for in vitro transcription followed by the colour code for every sequence. Start and stop codons are in bolt and underlined.

